# GLP-1R Agonism Directly Improves the Pumping Capacity of Murine Collecting Lymphatic Vessels

**DOI:** 10.64898/2025.12.26.696603

**Authors:** Mary E. Schulz, Victoria L. Akerstrom, Joseph H. Dayan, Jorge A. Castorena-Gonzalez

## Abstract

**Background:** Glucagon-like peptide-1 receptor (GLP-1R) agonists have recently been suggested as effective therapies to treat or reduce the risk of developing secondary lymphedema in patients with obesity; however, it is unknown whether the observed improvement in lymphatic function is solely due to weight loss-associated systemic benefits or in synergy with a lymphatic-specific effect of these pharmacological therapies.

**Methods:** We assessed the expression and localization GLP-1Rs in and around the lymphatic vasculature by single-cell RNA sequencing and fluorescence confocal microscopy. Using pressure myography we evaluated the direct effects of GLP-1R agonist, semaglutide, on modulating the contractile activity of lymphatic vessels from healthy wild-type (WT) mice, as well as lymphatics from diet-induced obese (DIO) WT mice, and hypercholesterolemic ApoE KO mice.

**Results:** Expression of *Glp1r* (encoding GLP-1Rs) was detected solely in LECs and was highly enriched in LECs from collecting lymphatics but absent in LECs from capillary regions. Pharmacological activation of GLP-1Rs using semaglutide led to robust vasodilation and an increase in the pumping capacity of isolated collecting lymphatics from WT, DIO, and ApoE KO mice. Compared to WT controls, lymphatics from ApoE KO mice displayed significant contractile dysfunction, which was restored with semaglutide. The GLP-1R-mediated response was in part facilitated by nitric oxide (NO), NADPH oxidase-mediated reactive oxygen species (ROS), and potentially vasodilatory prostanoids.

**Conclusions:** Our results revealed a direct, beneficial effect of GLP-1R agonism on lymphatic pumping capacity mediated by robust vasodilation, allowing lymphatics to accommodate larger fluid volumes, while maintaining strong and highly efficient contractions. Our observations implicated NO, ROS, and potentially vasodilatory prostanoids in the underlying mechanism; however, additional signaling components remain to be elucidated. These findings support recent clinical reports and further suggest that GLP-1R agonism could be an effective therapy for improving lymphatic contractile function in secondary lymphedema.

## Introduction

Lymphedema is a chronic, debilitating disease affecting >250 million people worldwide *with no pharmacological therapies available*^1–6^. In the U.S. and other developed countries, the largest group of patients afflicted with secondary lymphedema are cancer survivors, who develop this disease following surgery, chemotherapy, and radiotherapy, or as a result of cancer progression itself^7,8^. Depending on the type of malignancy, anatomical location, and type of treatments, the incidence of cancer-related lymphedema can vary largely^9,10^. For instance, the estimated incidence of breast cancer-related lymphedema is approximately 25%; however, multiple factors including high body mass index (BMI) and insulin resistance are known to significantly increase the risk (>3-fold) of developing this disease^11–18^. In fact, clinical studies have now demonstrated that obesity alone can cause secondary lymphedema^11,12,14,16,17,19–21^. Studies using animal models have shown that obesity and metabolic syndrome-induced lymphatic insufficiency is associated with impaired pumping capacity, hyperpermeability, and valve dysfunction in collecting lymphatics, impaired immune cell migration, downregulation of lymphatic endothelial cell (LEC) specific markers, and increased infiltration of immune cells and inflammatory molecules surrounding lymphatic networks among others^22–37^. The incidence of obesity-induced lymphedema continues to increase in association with the current obesity epidemic. A recent report by the Centers for Disease Control and Prevention (CDC) indicated that in the U.S., the prevalence of obesity (i.e., BMI≥30) in adults was 40.3%, and 1 in 10 adults were affected by severe obesity (i.e., BMI≥40)^38^. Therefore, there is a critical need to identify the molecular mechanisms of lymphatic dysfunction in obesity to develop the first pharmacological therapeutics to treat, prevent, or reduce the risk of developing secondary lymphedema.

GLP-1 receptor (GLP-1R) agonists, originally developed to help control blood glucose levels in type 2 diabetes, have gained popularity for their effectiveness to reduce bodyweight, and were recently linked to metabolic improvement in atherosclerosis and dyslipidemia^39–43^. Two recent publications by our collaborator and co-author Dr. Joseph Dayan have suggested that GLP-1R agonists may be an effective treatment for lymphedema and may reduce the risk of lymphedema in patients undergoing lymphadenectomy^44,45^. While improvement in lymphatic function in these patients could be associated with an overall improved systemic health, i.e., weight management, improved metabolic function, and glycemic control, the case report by Crowley et al. pointed to a potential synergistic effect with lymphatic-specific agonism of GLP-1Rs. That study^45^ reported on the case of a 44-year-old female patient who developed lymphedema following neoadjuvant chemotherapy, mastectomy, and axillary lymph node dissection and radiotherapy. This was a lean patient who, in association with adjuvant chemo and hormonal therapy, experienced a significant increase in bodyweight from 49.9 kg (BMI 19.2 kg/m^2^) to 66.3 kg (BMI 24.9 kg/m^2^), still in the healthy weight category. Her weight gain was resistant to diet and exercise intervention, and therefore, started receiving treatment with GLP-1R agonists. Initially with liraglutide, with limited weight loss response, and subsequently with semaglutide. Relevant to the central focus of this study, this patient lost 24% of her bodyweight in a period of 13 months, and more importantly, her lymphedema nearly completely resolved. An additional case report also demonstrated beneficial effects of GLP-1R agonism to reduce body weight, while improving fluid drainage and reducing limb circumference in a case of extreme obesity-induced massive, localized lymphedema^46^. The emergence of promising clinical evidence has prompted a formal prospective study of GLP-1R agonists in the treatment of lymphedema (personal communication, J. Dayan and colleagues December 2025). Of equal and critical importance is obtaining preclinical data to gain insight into the underlying mechanisms of how GLP-1 agonists may treat lymphatic dysfunction and lymphedema. However, hardly anything is known about the expression and distribution of GLP-1Rs in the lymphatic vasculature, let alone the roles that these receptors may play in the regulation of the function of lymphatic vessels. Two important questions surface which underpin the core of this paper: (1) are there GLP-1Rs in the lymphatic vasculature? and (2) if so, is there any observable effect on lymphatic function when these vessels are exposed to a GLP-1R agonist?

In this study, we first characterized the expression of GLP-1Rs and their distribution throughout the different cell types that make up the wall of collecting lymphatic vessels and surrounding tissues by means of a single-cell RNA sequencing (scRNAseq) dataset recently generated and published by our group^47^ and fluorescence confocal microscopy; then, we systematically assessed the specific effects of GLP-1R agonism on the contractile function of collecting lymphatic vessels from healthy C57BL/6J (WT) mice; and finally, we determined whether GLP-1R agonism could be employed to restore the contractile capacity of dysfunctional collecting lymphatics from diet-induced obese (DIO) and hypercholesterolemic ApoE KO mice, both models displaying multiple features commonly observed in metabolic syndrome.

## Methods

### Data Availability

Data and materials are publicly available in the main manuscript and within the Supplemental Materials. The referenced scRNAseq dataset was previously published by our group^37^ and was made available on NIH NCBI Gene Expression Omnibus (GEO accession: GSE294684).

### Animals

#### Mice

C57BL/6J (Cat. No.: 000664, WT) male and female mice and B6.129P2-Apoe^tm1Unc^/J (Cat. No.: 002052, ApoE KO) male mice were purchased from The Jackson Laboratory (Bar Harbor, ME) between 10-12 weeks old. Mice were kept in a temperature-controlled environment on a 12-hour light/dark cycle, with unrestricted access to standard food and water (Cat. No. 5053; PicoLab Rodent Diet 20 - Irradiated). Mice were given a minimum of 2 weeks before starting experiments for acclimatization. A subset of WT male and female mice were ordered at 4 weeks of age and fed a western diet (WD) for 16 weeks starting at 5 weeks of age (Cat. No. D12079Bi, Research Diets). All mice used in this study were between 3-6 months old. Mice were anesthetized with isoflurane, weighed, shaved, and prepared for terminal procedures and tissue dissection. All animal experiments were conducted in accordance with an approved IACUC protocol No.: 1884.

### Solutions and Chemicals

For dissection and cannulation of lymphatics, Krebs buffer was prepared with 146.9 mM NaCl, 5 mM D-glucose, 4.7 mM KCl, 3 mM NaHCO_3_, 2 mM CaCl_2_·2H_2_O, 1.5 mM Na-HEPES, 1.2 mM MgSO4, 1.2 mM NaH_2_PO_4_·2H_2_O, and 0.5% BSA (pH = 7.4). The buffer was sterile filtered and stored at 4°C. A separate Krebs buffer without BSA was used during pressure myography experiments for perfusion of the vessel chamber bath.

The following pharmacological agents were used for pressure myography experiments: GLP-1R agonist, semaglutide (1nM-1µM, Sema), L-NAME (100µM), indomethacin (10µM, Indo), and apocynin (100µM, Apo).

Chemicals and reagents were purchased from Sigma-Aldrich, TOCRIS, and MedChemExpress and these are listed in a Major Resources Table (Supplemental Materials).

### Microdissection of Collecting Lymphatic Vessels, and Pressure Myography

Inguinal axillary collecting lymphatic vessels were microdissected from the mouse as previously described^23,47–51^. A Sylgard coated dissection dish filled with BSA-containing Krebs buffer was used for pinning down tissues for fine dissection, where the majority of the adipose and connective tissues were removed. The lymphatic vessels were transferred into a custom-made pressure myography chamber with Krebs buffer, cannulated on glass micropipettes, pressurized equally at 3 cmH_2_O, and equilibrated with bath perfusion at 37°C for 30 minutes before experimentation. No-flow conditions were used in all pressure myography experiments (i.e., equal inflow and outflow pressures). For fine control of intraluminal pressures, OB1 MK3 microfluidic flow control systems (Elveflow, Paris) were used. Brightfield video recordings of the contractile activity of lymphatic vessels, as well as live tracking of inner and outer diameter changes were automatically recorded at 20 fps using custom-written Python-based programs.

### Contractile Parameters

Following experimentation, additional custom Python-based tools were used for the automated processing and analysis of contractile activity of lymphatic vessels (i.e., live-tracked changes in diameter as a function of time). As our previous studies^47–59^ have demonstrated, these analyses include calculating the mean values of contraction amplitude, end-diastolic diameter (EDD), end-systolic diameter (ESD), ejection fraction (EF), fractional pump flow (FPF), contraction frequency, and width of contraction. For this study, we have additionally calculated the average volume displaced per contraction, since the volume of fluid transported is an important metric for lymphatic function, particularly in lymphedema patients, where an improvement in volume transported would be extremely beneficial towards lessening fluid accumulation and increasing the amount of fluid shuttled back into the circulatory system. Volume displaced per contraction is determined via calculating the volume of the vessel (using the average length of a lymphangion (∼1mm), which is the length of the pumping unit of a lymphatic vessel) multiplied by the ejection fraction. Volume displaced is reported in nL of fluid. An important assumption when calculating volume displaced is valve competence, which allows for the generation of propulsive forward pressure.

#### Live Imaging of GLP-1R Expression in Isolated and Pressurized Collecting Lymphatics

The cellular localization of GLP-1Rs in collecting lymphatic vessels was assessed using semaglutide conjugated with FITC (Cat. No.: HY-114118F, MedChemExpress). Lymphatic vessels were dissected and prepared for cannulation as described above. A glass micropipette was pre-filled with 50µM semaglutide-FITC (dissolved in Krebs buffer) and the input end of the lymphatic vessel was cannulated onto that glass micropipette. Semaglutide-FITC was carefully flushed through the lumen by slowly increasing the pressure to ∼5 cmH_2_O. The vessel was then removed from the glass pipette, transferred into a dish containing Krebs-BSA buffer for rinsing. The vessel chamber was thoroughly rinsed, and the glass micropipette was washed to remove any residual semaglutide-FITC. The lymphatic vessel was then re-cannulated in a clean pressure myography chamber; the lumen was flushed to remove any unbound semaglutide-FITC still present within the lumen. The pressure myography chamber bath was exchanged with Ca^2+^-free Krebs, so the vessel did not develop spontaneous contractions during imaging. The lymphatic vessel was imaged at 25x magnification using a high-resolution and high-speed confocal imaging platform ANDOR Dragonfly 202 (Oxford Instruments) and a Zyla PLUS 4.2 Megapixel sCMOS camera. Images in three different regions of interest were collected from each lymphatic vessel that was stained.

### Single Cell RNA Sequencing Data Analysis

Analyses were performed using Seurat (4.3.0.1) in R studio (RStudio 2023.06.0+421). For our previously published scRNAseq dataset (see Data Availability section)^37^, we had performed the following quality control, cells with feature counts in the range 1000-6250 and cells displaying less than 5% mitochondrial counts were selected. Following quality control, the dataset underwent pipeline processing that included normalization using VST, multi-dimensional reduction, Harmony-based batch correction (with dims.use=1:30, representing the principal components to be utilized), cell clustering (with cluster resolution=0.8), differential testing framework, and visualization.

### Statistical Analyses

Results were analyzed with GraphPad Prism Version 10.6.1. Paired parametric t-test, unpaired parametric t-test with Welch’s correction (with no assumption of equal variance), two-way ANOVA corrected with Tukey test for multiple comparisons, and one-way ANOVA test that included a correction for multiple comparisons using Dunnett’s test and a Geisser-Greenhouse correction as no equal variances were assumed (i.e., sphericity was not assumed) were used to determine differences in contractile function parameters, with statistical significance set at p<0.05. Statistical tests were specified in the captions of each figure legend. For pressure myography experiments, the number “n” of experiments corresponds to each individually tested collecting lymphatic segment. Depending on the group or protocol, mice ranging from 6 to 14 were utilized; and frequently, multiple lymphatic segments were tested per mouse.

## Results

### Single-Cell Transcriptomic Characterization of Glp1r Expression in Collecting Lymphatic Vessels from WT Male and Female Mice

We previously performed single-cell RNA sequencing (scRNAseq) using collecting lymphatic vessels from male and female WT mice (n=4 mice per group). Briefly, we prepared single-cell suspensions from individually microdissected lymphatic vessels and surrounding connective tissues. From each mouse, collecting lymphatic vessels from multiple anatomical regions were pooled into a single sample (i.e., inguinal axillary (2 vessels), superficial cervical (4 vessels), popliteal afferent (4 vessels), and mesenteric (4 vessels)) in order to reach the minimum cell-count for scRNAseq experimentation. Automated clustering revealed 29 cell type clusters, and these were associated with 17 different cell types (Figure 1A). Unlike what has been reported in the blood vasculature, we found that *Glp1r* expression was exclusively detected in a subset of LECs (Figure 1A), with approximately 19.8±2.4% of LECs displaying detectable expression of *Glp1r*. Furthermore, there were no differences in *Glp1r* gene expression between lymphatics from female and male mice (Figure 1B). To investigate differences in *Glp1r*-expressing LECs vs. those that do not express this gene, we performed a differential gene expression (DGE) analysis. In LECs expressing *Glp1r*, DGE analysis identified a set of 10 differentially enriched (i.e., upregulated) genes, including Ptgs1, Fabp4, Ptn, Fabp5, Adamts9, Ndufa8, Naaa, Mir100hg, Lsr, and Nedd4l; while only 2 genes (i.e., Ccl21a and Sh3gl3) displayed significant downregulation (Figure 1C). Using the list of 10 differentially upregulated genes, a gene ontology (GO) analysis revealed several important molecular functions and biological processes involving fatty acid binding and transport, and prostanoid synthesis, in addition to roles in barrier integrity and metabolic processes (Figure 1D). To validate the presence of GLP-1R at protein level, we utilized FITC-labeled semaglutide to stain live lymphatic vessels and identify cellular localization of GLP-1Rs. Lymphatic vessels from n=4 WT mice were luminally flushed with 50 µM FITC-labeled semaglutide (see Methods section for a detailed description). Fluorescence confocal imaging confirmed that FITC-labeled semaglutide solely bound to LECs, as indicated by the longitudinal, flow-oriented pattern in the representative (from n=4) image in Figure 1E.

**Figure 1.**
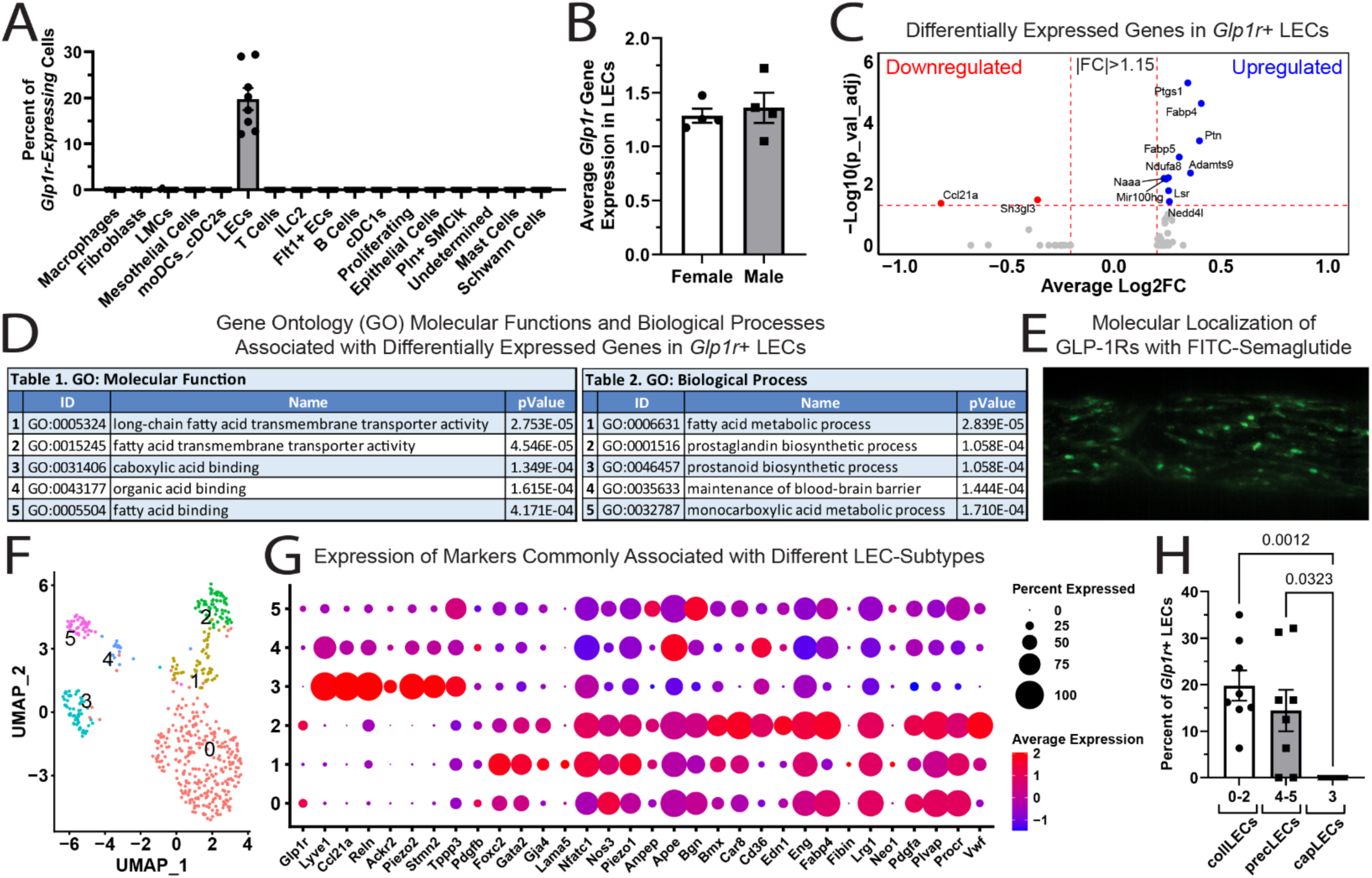
Expression of *Glp1r* in murine collecting lymphatic vessels. **(A)** Percent of *Glp1r*-expressing cells per cell identity; **(B)** Average gene expression of *Glp1r* in LECs from WT female and WT male mice respectively; **(C)** Volcano plot displaying the differentially expressed genes in LECs expressing *Glp1r* vs. those that did not express this gene; **(D)** Gene ontology analyses displaying the top molecular functions and biological processes associated with the differentially upregulated genes shown in panel C; **(E)** Representative image of an inguinal axillary lymphatic vessel exposed to FITC-labeled semaglutide to determine GLP-1R expression and cellular localization; **(F)** UMAP displaying the different LEC subtypes contained within the employed scRNAseq dataset; **(G)** DotPlot displaying the expression of *Glp1r*, as well as known markers of various LEC-subtypes (in this panel, the percent of cells expressing a given feature and the average gene expression are encoded in the size and color of dots respectively); and **(H)** Percent of LECs expressing *Glp1r* in clusters linked to collecting (collLECs), pre-collecting (precLECs), or capillary (capLECs) lymphatics (cluster numbers are indicated under the horizontal axis).

Using our scRNAseq dataset, we then performed subclustering of the LECs identity to determine the presence of different LEC-subtypes; our analysis identified 6 clusters (i.e., 0-5) with unique transcriptomic profiles (Figure 1F). Using markers previously reported to be differentially expressed in the different subtypes of LECs (e.g., capillary, collecting, pre-collecting, valve, etc.)^60^, we confirmed the presence of LECs from 1) Clusters 0-2: collecting lymphatics (including valve LECs); 2) Cluster 3: initial/capillary lymphatics; and 3) Clusters 4-5: precollecting lymphatics (Figure 1G). *Glp1r* expression was enriched in LECs from collecting and pre-collecting lymphatics (Figure 1G,H). Interestingly, *Glp1r* expression was not detected in LECs from the capillary region (Figure 1G,H). The expressions of *Foxc2*, *Gata2*, *Gja4*, and *Lama5* suggest that Cluster 1 may encompass LECs from lymphatic valves, which also displayed expression of *Glp1r*.

### Pharmacological Activation of GLP-1Rs by Semaglutide Modulates the Contractile Capacity of Collecting Lymphatic Vessels in a Concentration-Dependent Manner

Initially developed for the treatment of type 2 diabetes, GLP-1R agonists have been successful at treating obesity and improving weight loss, especially for patients whose obesity is resistant to diet and exercise alone. With the tight association between obesity worsening lymphatic dysfunction and lymphatic dysfunction worsening obesity, there is a critical need to assess the role of GLP-1R agonists in lymphatic function. Impaired pumping capacity of collecting lymphatic vessels contributes to abnormal interstitial fluid drainage and fluid accumulation. Therefore, we assessed the concentration-dependent (in the range 1-1000 nM) effects of the GLP-1R agonist semaglutide in isolated, cannulated, and pressurized collecting lymphatic vessels. Our results demonstrated that collecting lymphatic vessels are exquisitely sensitive to GLP-1R agonism, as indicative by the robust vasodilation observed even when vessels are exposed to 1 nM semaglutide (see representative diameter trace from a concentration response experiment in Figure 2A). Contraction amplitude was not affected by the increasing concentration of semaglutide (Figure 2B). Lymphatic vessels continued to vasodilate in a concentration-dependent manner, as shown by the increasing mean end diastolic and end systolic diameters (Figure 2C,D). Similar to contraction amplitude, mean ejection fraction and mean contraction width (at half-maximum) were also unchanged (Figure 2E,G). Contraction frequency and, therefore, the calculated fractional pump flow (FPF) were significantly reduced with each increasing concentration (Figure 2F,H). In this study, we incorporated a new calculated parameter that represents the mean volume displaced per contraction; this parameter was calculated for a 1-mm long lymphatic segment, which approximates the length of a lymphangion or pumping unit, and it is expressed in nL. Importantly, the mean volume displaced per contraction was significantly increased upon stimulation with semaglutide (Figure 2I). These results demonstrated that while GLP-1R agonism resulted in reduced contractile frequency, lymphatic vessels were capable of accommodating larger fluid volumes (i.e., vasodilated) and displayed strong and more efficient contractions displacing larger fluid volumes.

**Figure 2.**
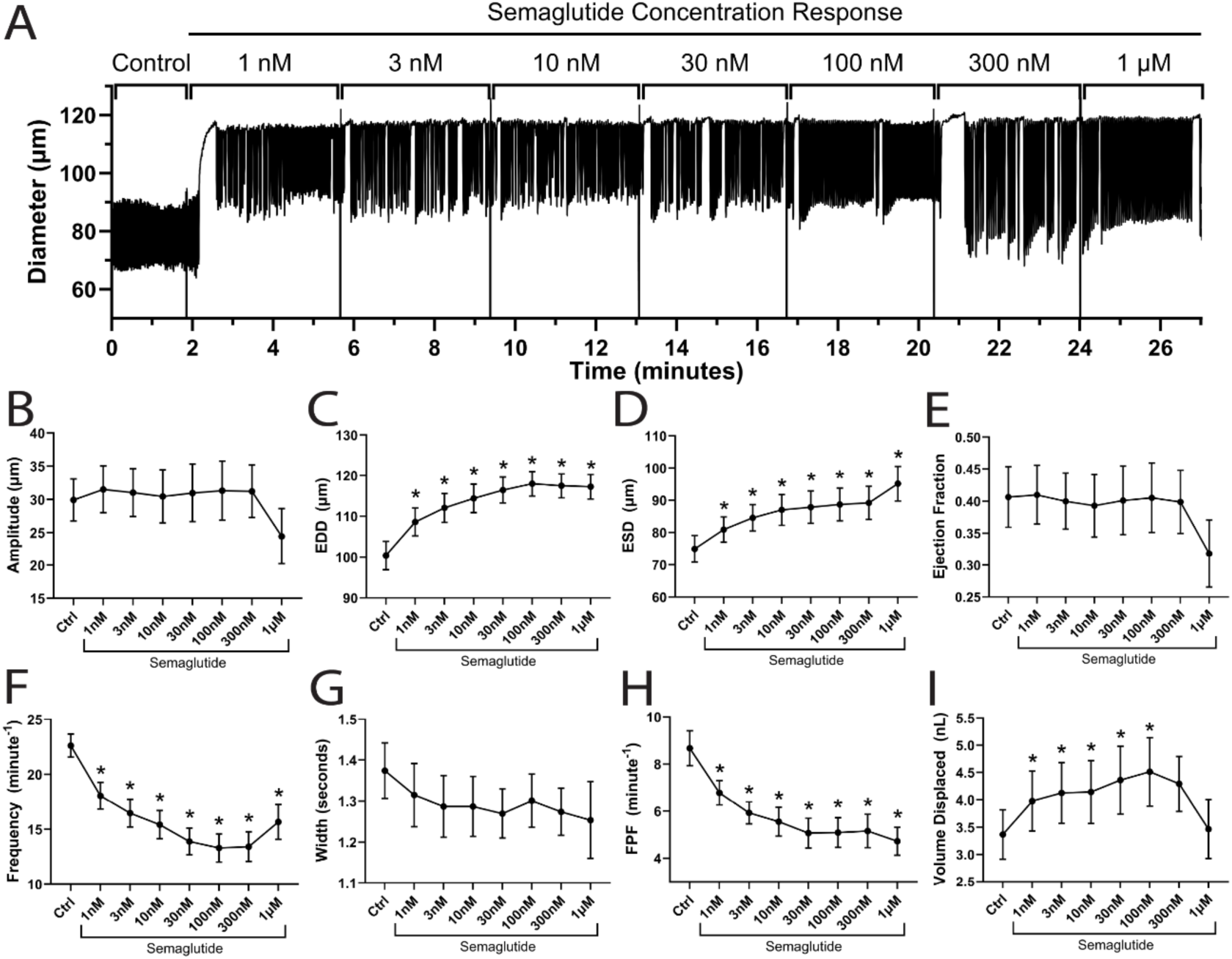
Concentration response curve of GLP-1R agonist, semaglutide on lymphatic vessels from WT mice. **(A)** Representative diameter trace of an inguinal axillary lymphatic vessel displaying the functional response to increasing concentrations of semaglutide in the range of 1nM to 1µM; the contractile activity was assessed for 5 minutes at each concentration; **(B-I)** Mean contractile parameters as a function of semaglutide concentration (expressed as mean±SEM), including **(B)** amplitude, **(C)** end diastolic diameter (EDD), **(D)** end systolic diameter (ESD), **(E)** ejection fraction, **(F)** contraction frequency, **(G)** width, **(H)** fractional pump flow (FPF), and **(I)** volume displaced. A total of n=17 vessels were included and a one-way ANOVA test that included a correction for multiple comparisons using Dunnett’s test and a Geisser-Greenhouse correction as no equal variances were assumed (i.e., sphericity was not assumed) were used to determine differences in contractile function parameters, with statistical significance set at p<0.05.

### Acute Treatment with Semaglutide Improves Pumping Capacity of WT Collecting Lymphatic Vessels

In the previous section, our data demonstrated that the contractile activity of lymphatic vessels was significantly modulated by semaglutide at concentrations as low as 1 nM; and we determined that the maximal beneficial effects (i.e., maximum vasodilation and volume displaced), while maintaining rhythmic contractility, were achieved at concentrations in the range of 1-10 nM. Therefore, we examined and characterized the functional responses in WT inguinal axillary lymphatic vessels following a single acute stimulus with 5 nM semaglutide (while constant superfusion with Krebs buffer was maintained). The contractile activity was recorded and analyzed for 2 minutes under control conditions and following stimulation with semaglutide. A representative trace from a total of n=11 is shown in Figure 3A. Consistent with our concentration response experiments, following a single, acute dose with 5 nM semaglutide, the contraction amplitude of lymphatics was not significantly changed, although there was a trend for amplitude to be increased after treatment (Figure 3B); lymphatic vessels displayed robust vasodilation as indicated by the increased end diastolic (EDD) and end systolic (ESD) diameters (EDD 108.9±6.8µm (Ctrl) vs. 124.3±8.0µm (Sema), p<0.0001, and ESD 62.95±4.3µm (Ctrl) vs. 74.53±5.8µm (Sema), p<0.0032)(Figure 3C,D); ejection fraction and contraction width (at half-maximum) remained unchanged (Figure 3E,G); contraction frequency was significantly decreased after GLP-1R agonism with semaglutide (i.e., 19.23±0.98 contractions/minute (Ctrl) vs. 12.13±0.84 contractions/minute (Sema)) and so did the calculated fractional pump flow (Figure 3F,H); however, the calculated volume displaced by each contraction (Figure 3I) was significantly increased after a single 5 nM stimulus with semaglutide (i.e., 6.91±1.70nL (Ctrl) vs. 8.63±2.03nL (Sema), p<0.001). To further exemplify this increase in pumping capacity, Figure 3J depicts the calculated volume displaced for each individual contraction in the representative trace being shown in Figure 3A. Note that in the initial 2-minutes under control conditions, each contraction displaced ∼7.5nL, and this was significantly increased to >10nL after stimulation with semaglutide. Mean values were then calculated for all contractions within the control period, as well as following treatment with semaglutide, and these are represented as a single datapoint in Figure 3I.

**Figure 3.**
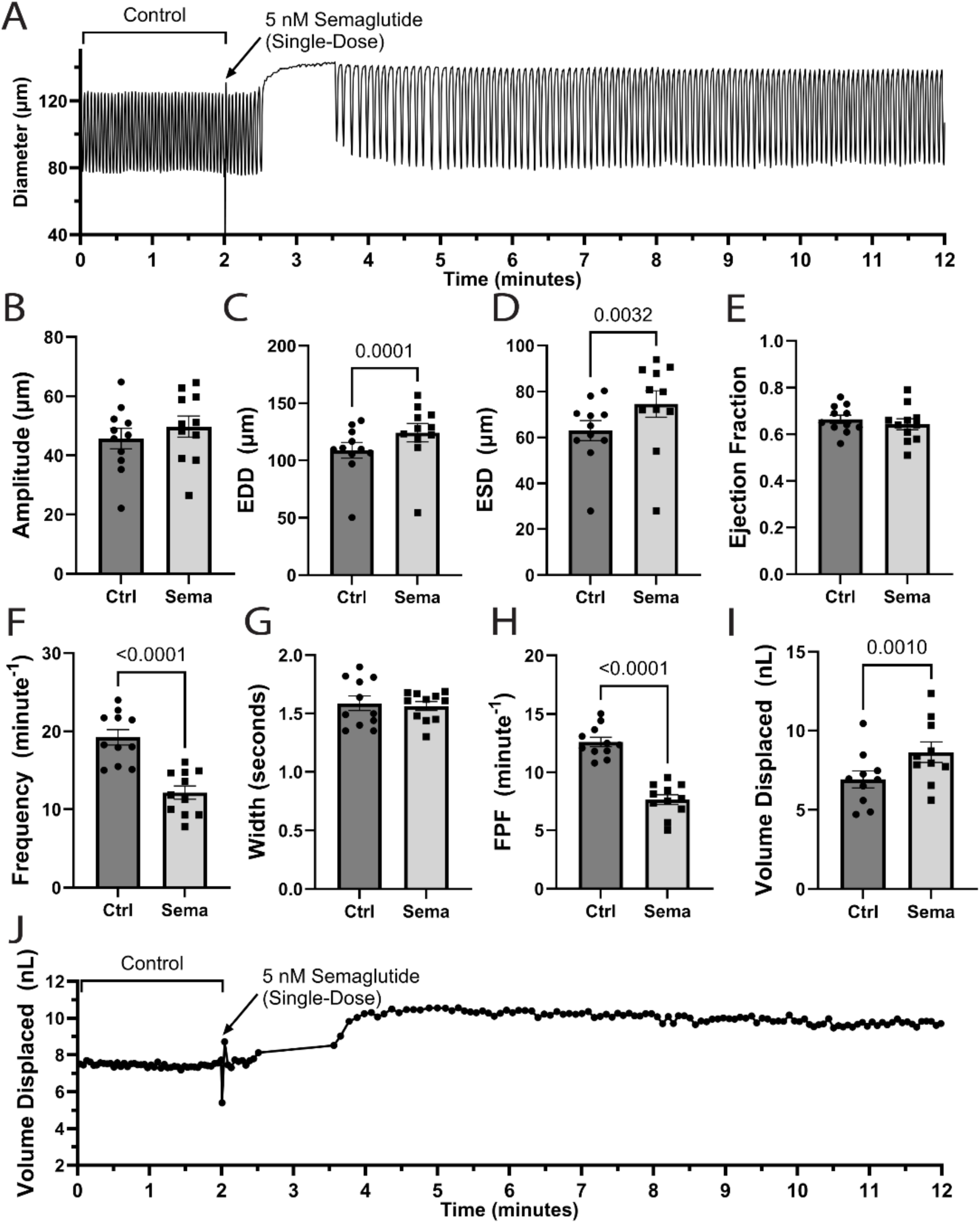
Acute stimulation with a 5 nM semaglutide single dose improves the pumping capacity of lymphatic vessels. **(A)** Representative diameter trace displaying the contractile activity of an inguinal axillary lymphatic vessel from a WT mouse under control conditions and following stimulation with a single 5 nM dose of semaglutide. **(B-I)** Summary data of different contractile parameters before (Ctrl) and after 5nM stimulus with semaglutide (Sema), including **(B)** amplitude, **(C)** end diastolic diameter (EDD), **(D)** end systolic diameter (ESD), **(E)** ejection fraction, **(F)** contraction frequency, **(G)** width, **(H)** fractional pump flow (FPF), **(I)** volume displaced, and **(J)** volume displaced for each individual contraction in panel A. A total of n=11 vessels were used for these experiments and paired parametric T test was used to determine significance, with statistical significance set at p<0.05.

### Semaglutide-Induced Increase in the Pumping Capacity of Collecting Lymphatics is Only Partially Mediated by LEC-Derived Nitric Oxide and Vasodilatory Prostanoids

To determine the mechanism by which semaglutide improves lymphatic pumping capacity and to determine the potential, suspected involvement of nitric oxide (NO) and vasodilatory prostanoids, we assessed the functional responses of collecting lymphatic vessels from WT mice to 5 nM semaglutide following a 20-minute pre-treatment with L-NAME (100µM) and indomethacin (10µM), which inhibit nitric oxide synthase (NOS) and cyclooxygenases-1,2 (COX-1 and COX-2) respectively. Compared to control conditions, superfusion with a Krebs buffer containing L-NAME and indomethacin resulted in an increase in basal vessel tone, as indicative by the reduced EDD and ESD (Figure 4C,D), an increase in ejection fraction and contraction width; while contraction amplitude, frequency, fractional pump flow (FPF), and volume displaced remained unchanged (Figure 4B,E-I). In the presence of L-NAME and indomethacin, semaglutide (5nM) still induced significant changes in lymphatic contractility, including increased contraction amplitude (Figure 4B), partial but significant vasodilation (Figure 4C,D), increased contraction width (Figure 4G), and decreased frequency and FPF (Figure 4F,H). The latter observation demonstrating a significant decrease in contraction frequency following treatment with semaglutide and despite the presence of L-NAME and indomethacin (i.e., expressed in contractions/minute 20.87±0.69 (Ctrl) vs. 19.64±0.73 (L-NAME+Indo) vs. 17.41±0.68 (L-NAME+Indo+Sema), p<0.01), indicates that the semaglutide-induced decrease in contractile frequency was not solely due to the vasodilatory actions of NO and/or the potential secretion of prostanoids (Figure 4F). Furthermore, while the calculated volume displaced per contraction after treatment with semaglutide was significantly increased from that in the presence of L-NAME and indomethacin, this was not significantly different from that under control conditions (Figure 4I), suggesting a partial but necessary contribution of NO and/or potentially secreted prostanoids in the semaglutide-induced increase in the pumping capacity of collecting lymphatics. This data also points to additional NO- and prostanoid-independent signaling by which GLP-1R agonism modulates the contractile function of collecting lymphatic vessels.

**Figure 4.**
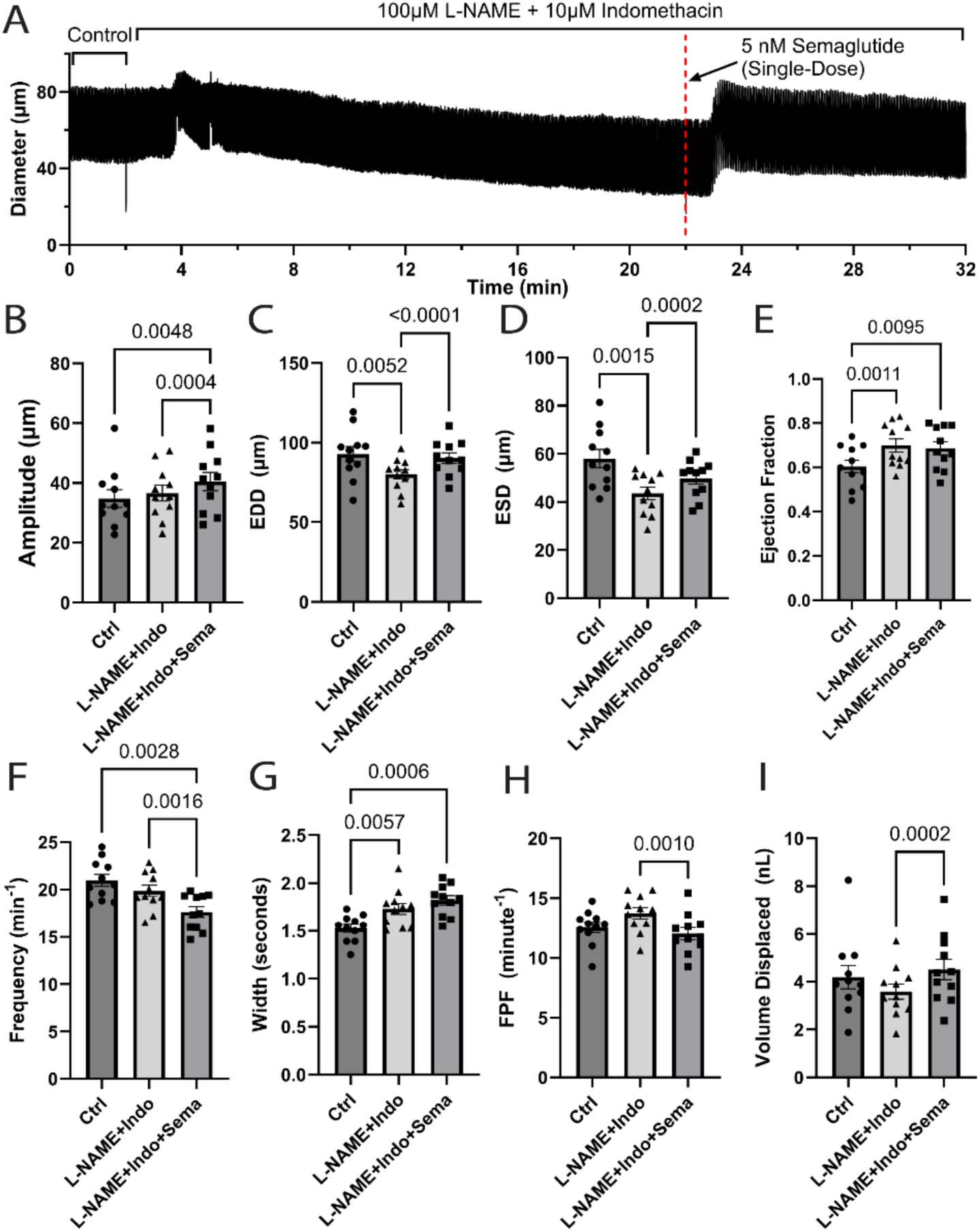
Role of nitric oxide and prostanoids in semaglutide-mediated modulation of lymphatic contractile function. **(A)** Representative diameter trace displaying the contractile activity of an inguinal axillary lymphatic vessel from a WT mouse under control conditions, during a 20-minute pre-treatment with L-NAME (100µM) and indomethacin (10µM), and after stimulation with a single 5 nM dose of semaglutide. **(B-I)** Summary data of different contractile parameters under these 3 paired conditions, including **(B)** amplitude, **(C)** end diastolic diameter (EDD), **(D)** end systolic diameter (ESD), **(E)** ejection fraction, **(F)** contraction frequency, **(G)** width, **(H)** fractional pump flow (FPF), and **(I)** volume displaced. A total of n=11 vessels from WT mice were used for these experiments. A one-way ANOVA corrected with a Geisser-Greenhouse correction as no equal variances were assumed (i.e., sphericity was not assumed) was used to determine differences in contractile function parameters, with statistical significance set at p<0.05.

### NADPH Oxidase-Mediated Reactive Oxygen Species Contribute to the Basal Regulation of Collecting Lymphatic Vessel Contractility and May Be Involved in the Semaglutide-Induced Increase in Lymphatic Pumping Capacity

Our results presented in the previous section suggested that while NO and potentially secreted prostanoids may play a crucial role in mediating the semaglutide-induced lymphatic vasodilation, while maintaining strong, and in fact, more efficient contractions; these results also pointed to the presence of other signaling, perhaps other paracrine signaling originating from the lymphatic endothelium and reaching lymphatic muscle cells (LMCs). Reactive oxygen species (ROS) are known to play critical roles in vascular physiology, including vasodilation, in health and disease. Therefore, to determine the potential contribution of major ROS, including hydrogen peroxide, we performed additional experiments on isolated collecting lymphatic vessels exposed to 5 nM semaglutide (single dose) following a 20-minute pre-treatment and in the presences of L-NAME (100µM), indomethacin (10µM), and the NADPH oxidase inhibitor apocynin (100µM). In contrast to the functional responses shown in Figure 4, where L-NAME+Indo induced a significant increase in ejection fraction, while contraction amplitude and volume displaced remained unchanged (Figure 4), the addition of apocynin (i.e., L-NAME+Indo+Apo) resulted in a significant decrease in contraction amplitude and volume displaced (Figure 5A,I), with no changes in ejection fraction. Pre-treatment with either L-NAME+Indo or L-NAME+Indo+Apo led to an increase in tone, i.e., significantly decreased EDD when compared to control conditions (i.e., Ctrl); however, the decrease in EDD induced by L-NAME+Indo+Apo (16.3±3.8µm) was significantly higher than that induced by L-NAME+Indo (12.5±3.0µm)(Figure 4C and Figure 5C). These results suggest a potential contribution of NADPH oxidase in regulating basal lymphatic contractility. In the presence of L-NAME+Indo+Apo, semaglutide still induced a significant increase in contraction amplitude, EDD, and volume displaced; however, these values were still comparable or below those observed in control conditions (Figure 5B,C,I). These results demonstrate that while the increase in pumping capacity induced by semaglutide is prevented in the presence of L-NAME+Indo or L-NAME+Indo+Apo, the residual response observed in lymphatic vessels following stimulation with semaglutide even in the presence of these inhibitors points to additional, yet to be identified, signaling contributions.

**Figure 5.**
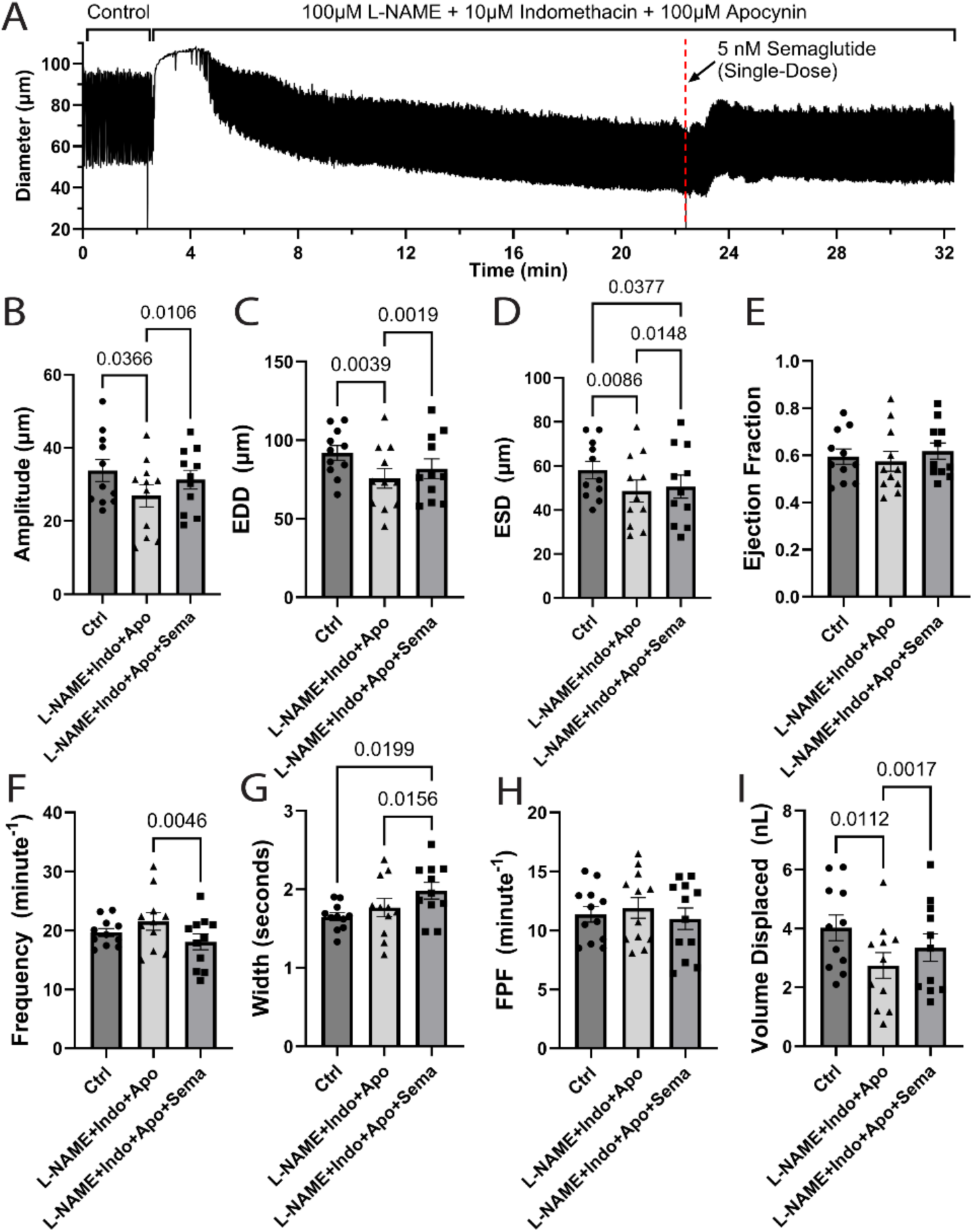
Role of NADPH oxidase-mediated ROS in semaglutide-mediated modulation of lymphatic contractile function. **(A)** Representative diameter trace displaying the contractile activity of an inguinal axillary lymphatic vessel from a WT mouse under control conditions, during a 20-minute pre-treatment with L-NAME (100µM), indomethacin (10µM), and apocynin (100µM), and after stimulation with a single 5 nM dose of semaglutide. **(B-I)** Summary data of different contractile parameters under these 3 paired conditions, including **(B)** amplitude, **(C)** end diastolic diameter (EDD), **(D)** end systolic diameter (ESD), **(E)** ejection fraction, **(F)** contraction frequency, **(G)** width, **(H)** fractional pump flow (FPF), and **(I)** volume displaced. A total of n=11 vessels from WT mice were used for these experiments. A one-way ANOVA corrected with a Geisser-Greenhouse correction as no equal variances were assumed (i.e., sphericity was not assumed) was used to determine differences in contractile function parameters, with statistical significance set at p<0.05.

### Semaglutide Improves the Pumping Capacity of Lymphatic Vessels from Diet-Induced Obese (DIO) WT and Hypercholesterolemic ApoE KO Mice

In previous sections, we characterized the effects of a single stimulus of semaglutide to the contractile function of healthy collecting lymphatics from WT mice, while the lymphatic vessels were maintained under constant superfusion with fresh Krebs buffer, allowing for drug washout. We then aimed to 1) characterize the functional response in collecting lymphatics when the concentration of semaglutide was maintained for a longer period of time, and 2) determine the potential beneficial effects of semaglutide to improve the contractile capacity of inguinal axillary lymphatic vessels from diet-induced obese (DIO) and ApoE KO mice. These are both animal models that recapitulate various components of metabolic syndrome and display various aspects of lymphatic system dysfunction. Following a 30-minute equilibration, the contractile activity of lymphatic vessels was first recorded under control conditions, for 2 minutes, and then the concentration of semaglutide in the bath was increased to 5 nM, and the functional response of collecting lymphatics was recorded for 30 minutes. The concentration of semaglutide was maintained by constant superfusion with a Krebs buffer containing the same concentration of the drug (i.e. 5 nM semaglutide). In control conditions and compared to lymphatics from WT mice, lymphatic vessels from DIO mice displayed significantly smaller diameters (Figure 6B,C); however, their contractile function was not significantly impaired (although contraction amplitude and volume displaced trended lower, Figure 6A,H). In contrast, lymphatic vessels from ApoE KO mice had similar basal diameters when compared to WT controls but displayed significantly decreased contraction amplitude (31.45±1.47µm for WT vs. 21.73±2.39µm for ApoE KO), ejection fraction (0.47±0.02 for WT vs. 0.34±0.03 for ApoE KO), and volume displaced (4.96±0.27nL for WT vs. 3.37±0.41nL for ApoE KO) (Figure 6A,D,H); while contraction frequency was significantly higher (Figure 6E). Consistent with our observations reported in the previous sections, the modulation of the contractile function by semaglutide occurred within the first couple minutes of exposure to the drug and these effects remained stable throughout the duration of the recording.

**Figure 6.**
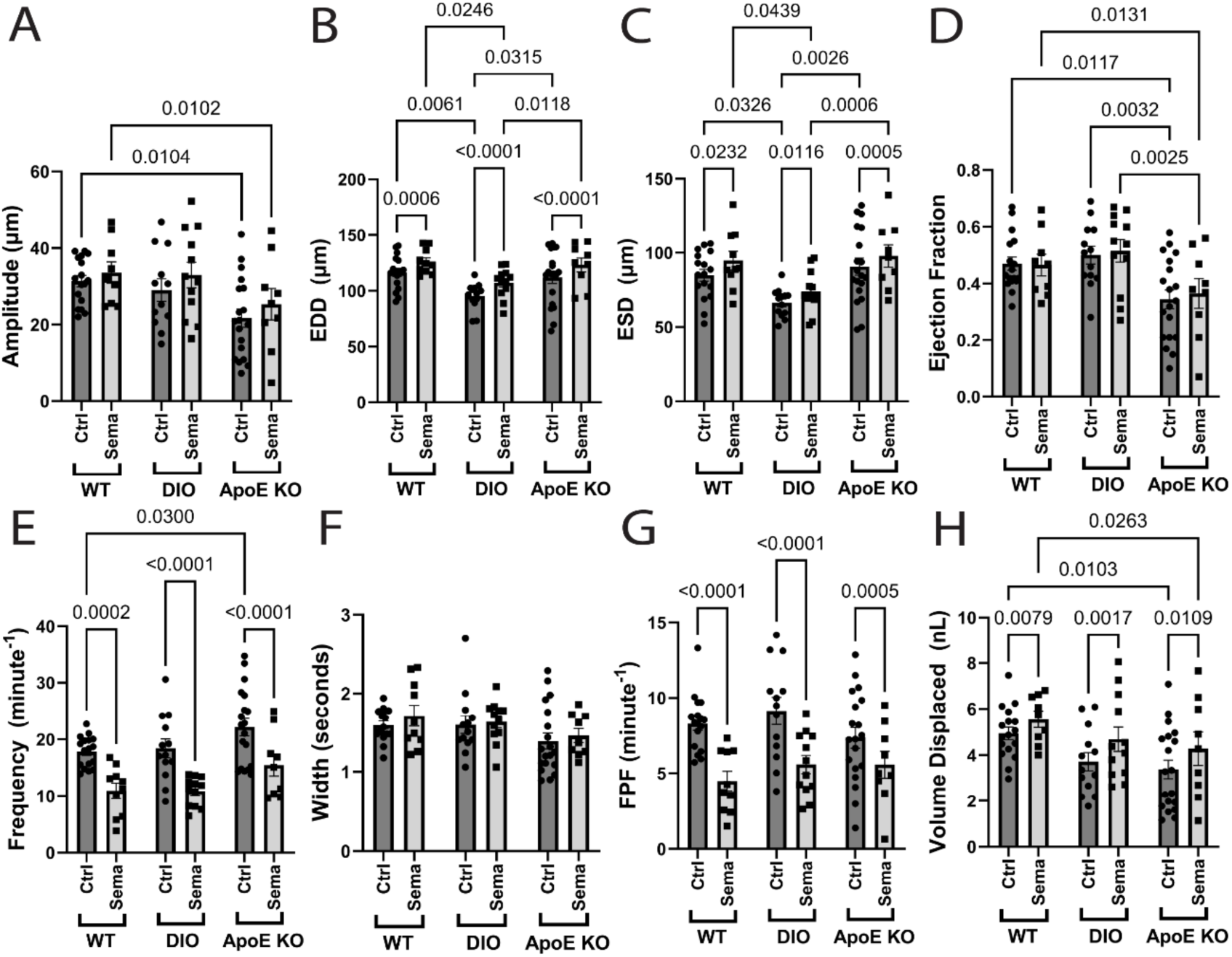
Semaglutide improves and restores the pumping capacity of dysfunctional lymphatic vessels from diet-induced obese (DIO) and hypercholesterolemic ApoE KO mice. Summary data of contractile parameters characterizing the contractile function of lymphatic vessels from WT, DIO, and ApoE KO mice under control conditions and following perfusion with a Krebs buffer containing 5 nM semaglutide. These contractile parameters are expressed as mean±SEM and include **(A)** contraction amplitude, **(B)** end diastolic diameter (EDD), **(C)** end systolic diameter (ESD), **(D)** ejection fraction, **(E)** contraction frequency, **(F)** contraction width (at half maximum), **(G)** fractional pump flow (FPF), and **(H)** volume displaced. A total of n=17 (control) and n=10 (sema) vessels from WT mice, n=13 (control) and n=12 (sema) vessels for DIO mice, and n=19 (control) and n=9 (sema) vessels from ApoE KO mice were used for these experiments. A two-way ANOVA was utilized to assess differences in contractile parameters with statistical significance set as p<0.05.

Importantly, the impaired contractile amplitude, ejection fraction, and volume displaced per contraction observed in lymphatics from ApoE KO mice were all improved and restored to values comparable to those of lymphatics from WT mice in control conditions by superfusion with semaglutide (Figure 6A,D,H). In fact, volume displaced, a parameter that estimates fluid transport per contraction, was significantly increased with semaglutide perfusion in lymphatics from all groups, i.e., 4.96±0.27nL (Ctrl) vs. 5.56±0.34nL (Sema) for WT, 3.72±0.40nL (Ctrl) vs. 4.70±0.52nL (Sema) for DIO, and 3.37±0.41nL (Ctrl) vs. 4.28±0.72nL (Sema) for ApoE KO (Figure 6H). Irrespective of the group, semaglutide perfusion induced robust vasodilation as indicative by the significant increases in EDD, i.e., 116.47±3.59µm (Ctrl) vs. 126.06±3.49µm (Sema) for WT, 95.35±3.30µm (Ctrl) vs. 107.19±3.82µm (Sema) for DIO, and 112.11±5.58µm (Ctrl) vs. 123.18±6.30µm (Sema) for ApoE KO (Figure 6B). Consistent with our previous observations, semaglutide induced a significant reduction in contractile frequency and calculated FPF (Figure 6E,G); while contraction width (at half-maximum) remained unaffected by group and semaglutide treatment (Figure 6F).

## Discussion

Cancer and obesity-related lymphedema make up the largest groups of patients afflicted with this disease, with no pharmacological therapies available. In recent years, GLP-1R agonists, initially developed and approved for treatment of type 2 diabetes, have been shown to be effective for fighting obesity and promoting weight-loss in patients who have adipose tissue that is refractory to diet and exercise alone. A few case studies have reported beneficial effects of GLP-1R agonists on the lymphatic system, specifically in improving lymphedema in patients with various degrees of obesity^45,46^ and in reducing the risk of developing secondary lymphedema in patients who underwent axillary lymph node dissection for breast cancer treatment^44^. However, prior to this study, the expression of GLP-1Rs in the lymphatic vasculature and the direct effects of GLP-1R agonists on the function of lymphatic vessels remained unexplored.

The lymphatic system relies on spontaneous, strong, and highly entrained contractions^48^ in order to transport lymph from the interstitial spaces back into the central circulation. Also critical are intraluminal valves, which not only ensure net unidirectional, forward flow, but they also allow for the generation of propulsive pressure during each contraction. Assessment of lymphatic contractile activity in patients with lymphedema is limited; however, a few studies have reported on decreased pumping capacity of collecting lymphatics in the affected lymphedematous limbs^61,62^. Therefore, as the lymphatic field continues to move forward and novel therapies emerge, it is crucial to consider the development of pharmacological alternatives that target the improvement and/or restoring of the pumping capacity of lymphatic vessels.

In this study, we aimed to first characterize the expression of GLP-1Rs in the lymphatic vasculature and, because of its importance in fluid drainage and transport, also evaluate the specific effects of GLP-1R agonism to the regulation of lymphatic contractile function.

Our scRNAseq observations showed that in a dataset that included lymphatic vessels and nearby surrounding tissues, expression of *Glp1r*, encoding GLP-1Rs, was solely detected in LECs. Furthermore, subsequent subclustering of LECs led to the identification of 6 unique LEC subtypes. We utilized markers previously reported by Dr. Tatiana Petrova’s group^60^ to annotate our LEC scRNAseq data and determined that these subclusters displayed transcriptomic profiles consistent with collecting LECs (including valve LECs), pre-collecting LECs, and capillary LECs. Intriguingly, expression of *Glp1r* was highly enriched in LECs from collecting and pre-collecting LECs but was not detectable in LECs from capillary/initial lymphatics.

We then utilized pressure myography on isolated, cannulated, and pressurized collecting lymphatics, a technique that allows us to study the ex-vivo function of lymphatic vessels under controlled conditions. Our results demonstrated, for the first time, a direct beneficial effect of GLP-1R agonism to the contractile function of lymphatic vessels. Even at the lowest tested concentration (i.e., 1 nM), semaglutide induced robust vasodilation, usually within the 30-60 seconds following exposure to the drug, suggesting that lymphatic vessels are likely sensitive to semaglutide at concentrations even <1 nM. The vasodilatory effects of semaglutide were maximal at concentrations in the range of 1-10 nM; in fact, a single stimulus with 5 nM semaglutide induced a sustained vasodilatory effect that lasted >10 minutes, even while the preparation was maintained under constant bath perfusion (∼0.2 mL/min). Other potent vasodilators induce robust vasodilation with a concomitant concentration-dependent loss of contractility, including contraction frequency and amplitude. Semaglutide also induced a significant reduction in contractile frequency; however, lymphatic vessels maintained strong contractions even at the higher range of concentrations included in this study (i.e., 1 µM). The significant increase in vessel diameter allowed lymphatic vessels to accommodate larger volumes of fluid resulting in more efficient pumping, as indicated by the significantly increased volume displaced, a new calculated parameter we have introduced in this study to estimate the amount of fluid that would be pumped/transported by a 1-mm long lymphangion (i.e., the lymphatic pumping unit).

Importantly, despite semaglutide decreasing contractile frequency, the calculated volume of fluid displaced with each contraction was significantly increased in lymphatic vessels from animal models that recapitulate various aspects present in metabolic syndrome (i.e., DIO and chow fed ApoE KO mice). Previously, studies led by Dr. Véronique Angeli and Dr. Gwendalyn Randolph demonstrated that ApoE KO fed a high-fat diet displayed lymphatic vessel degeneration, as well as multiple aspects of lymphatic dysfunction, including severe valve dysfunction and lymphatic contractile impairment^26,63^. In this study, our data demonstrated that lymphatic contractile dysfunction is present even in ApoE KO mice fed a regular chow, suggesting that hypercholesterolemia may be sufficient to induce contractile impairment. In fact, our preliminary unpublished observations suggest that lymphatic contractile function in chow-fed ApoE KO mice declines with age. Our lab is currently conducting additional studies, beyond the scope of this manuscript, to characterize contractile dysregulation in control chow-fed and western diet-fed ApoE KO mice to identify the underlying molecular mechanisms leading to severe dysfunction of collecting lymphatic vessels in obesity and hypercholesterolemia, including contractile and valve dysfunction, as well as hyperpermeability.

The strong vasodilatory effects of semaglutide were found to be partially linked to nitric oxide, potentially to secreted prostanoids, and NADPH oxidase-mediated ROS; however, a residual contractile response was consistently present even in the presence of pharmacological inhibitors of NOS, COX-1,2, and NADPH oxidase. These results suggest additional signaling, potentially of paracrine nature, yet to be elucidated. Alternatively, the unexplored possibility is that GLP-1Rs are present in LMCs (and/or other cell types) with low mRNA levels, beyond the level of detection in the experimental and sequencing conditions of the studies hereby included. In addition to endothelial expression, GLP-1R expression has been reported in various types of smooth muscle, including vascular smooth muscle, and their activation has been linked to vasodilation and increase in blood flow^64–71^. Our group attempted to validate the expression with and without localization of GLP-1R at a protein level using immunofluorescence and western blotting respectively; however, no usable data was obtained after testing numerous antibodies. Therefore, we employed FITC-labeled semaglutide, where no LMC pattern was detected in collecting lymphatics. Additional more in-depth validations are still required.

Once lymph enters the initial lymphatics, some of the key factors that determine efficient lymph transport along the collecting lymphatic vasculature are 1) efficient contractility, 2) the ability of collecting lymphatics to properly vasodilate as part of the contraction cycle, 3) competent intraluminal valves, and 4) a relative low permeability that ensures lymph remains within the lymphatic vasculature. All these factors are equally important and necessary to ensure efficient pumping of fluid and are likely impaired in lymphedema. NO and other paracrine vasodilators are not required for the relaxation of LMCs at the end of each contraction cycle; however, these do play an important role in determining the maximum systolic diameter that lymphatic vessels can reach during this relaxation. In this study, our observations are in agreement with recent clinical studies and support the therapeutic potential of semaglutide to improve lymphatic function in secondary lymphedema including obesity-related and cancer-related lymphedema.

Importantly, the exquisite sensitivity of collecting lymphatic vessels to GLP-1R agonism points to semaglutide, and other GLP-1R agonists, as effective therapies to improve lymphatic function independently of their weight-loss effects. The specific lymph concentrations of semaglutide in humans are unknown, and these likely vary between visceral and peripherally located lymphatics; however, direct uptake of GLP-1Rs by the lymphatic system has previously been demonstrated by Dr. Natalie Trevaskis’s group in animal models^72^. Upcoming work by Dr. Trevaskis will soon determine the specific concentrations of semaglutide in lymph. Our results suggest that significantly lower dosages of semaglutide, compared to those commonly prescribed for weight loss, could be employed to restore the pumping capacity of dysfunctional lymphatics in lymphedema. Low dosages (e.g., <100 nM) of semaglutide could be directly delivered (i.e., subdermal) into the affected limb minimizing its systemic side effects. While these speculations remain to be tested and validated, unpublished clinical studies presently being conducted by Dr. Joseph Dayan (shared with permission) have shown that 8 out of 9 patients in a prospective study on GLP-1R treatment for lymphedema have experienced a significant reduction in limb volume (∼10% average reduction) and improvement in validated quality of life scores (LLIS) by 36% with the affected limb having a higher volume reduction than the unaffected limb. Noteworthy, these beneficial effects of GLP-1R agonism were observed even after only 3 months of treatment with semaglutide at the lowest prescribed dosage of 0.25 mg/week.

In conclusion, our results specifically demonstrate a direct beneficial effect of semaglutide to improve fluid transport by allowing lymphatics to robustly vasodilate while maintaining efficient pumping activity; however, our findings also demonstrated that GLP-1Rs are highly enriched in LECs, where these channels may play additional important roles, even some potentially regulating lymphatic valve and barrier function (i.e., permeability), which are being investigated in separate studies from our group.

## Perspectives

Secondary lymphedema is most commonly associated with 1) cancer-related therapeutic interventions, including chemotherapy, mastectomy, axillary lymph node dissection, and radiotherapy, as well as with 2) obesity and metabolic syndrome, with no pharmacological therapies available. In agreement with recent clinical studies suggesting that pharmacological agonism of GLP-1Rs may be an effective therapeutic strategy to treat secondary lymphedema in these groups of patients, using murine models, this study demonstrated that GLP-1Rs are expressed in the lymphatic vasculature and are specifically enriched in the lymphatic endothelium. Furthermore, collecting lymphatic vessels from healthy mice, as well as from mouse models of obesity and metabolic syndrome displayed an exquisite sensitivity to pharmacological activation with semaglutide, which significantly increased or restored their pumping capacity. In lymphedema, impaired contractile/pumping function of lymphatic vessels is a known key determinant in the development and progression of the disease, the observations hereby presented, supported by clinical reports, suggest the therapeutic potential of GLP-1R agonists to directly restore the pumping capacity of collecting lymphatic vessels in secondary lymphedema.

## Abbreviations

Name: Abbreviation
GLP-1R: Glucagon-like Peptide-1 Receptor
scRNAseq: Single-Cell RNA Sequencing
PSS: Physiological Saline Solution
L-NAME: NG-Nitro-L-Arginine Methyl Ester
Indomethacin: Indo
Apocynin: Apo
Semaglutide: Sema
LECs: Lymphatic Endothelial Cells
LMCs: Lymphatic Muscle Cells
WT: Wild Type
DIO: Diet-Induced Obesity
WD: Western Diet

