## Supplemental Materials for "GLP-1R Agonism Directly Improves the Pumping Capacity of Murine Collecting Lymphatic Vessels"

### Major Resources Table

### Animals

| Species | Vendor or Source | Background Strain | Sex | Persistent ID / URL |
|---|---|---|---|---|
| Mouse | Jackson Lab | C57BL/6J (Wild-type, WT) | M & F | https://www.jax.org/strain/000664 |
| Mouse | Jackson Lab | B6.129P2-Apoe$^{tm1Unc}$/J (ApoE KO) | M | https://www.jax.org/strain/002052 |

### Reagents for Live Vessel Staining

| Target antigen | Vendor | Catalog # | Working concentration | Persistent ID / URL |
|---|---|---|---|---|
| Semaglutide-FITC | MedChemExpress | Hy-114118F | 50µM Ing ax lymphatic n=4 | https://www.medchemexpress.com/semaglutide-fitc-labeled.html |

### Data & Code Availability

| Description | Source / Repository | Persistent ID / URL |
|---|---|---|
| scRNAseq (GEO accession #: GSE294684) | NIH NCBI GEO | https://www.ncbi.nlm.nih.gov/geo/query/acc.cgi?acc=GSE294684 |

### Pharmacological Reagents for Pressure Myography Experiments

| Description | Source | Catalog # | Persistent ID / URL |
|---|---|---|---|
| Apocynin | Tocris | 4663 | https://www.tocris.com/products/apocynin_4663 |
| L-Name | Sigma | N5751 | https://www.sigmaaldrich.com/US/en/product/sigma/n5751 |
| Indomethacin | Tocris | 1708 | https://www.tocris.com/products/indomethacin_1708 |
| Semaglutide | MedChemExpress | HY-114118 | https://www.medchemexpress.com/Semaglutide.html |
| Semaglutide-FITC | MedChemExpress | HY-114118F | https://www.medchemexpress.com/semaglutide-fitc-labeled.html |

### Chemicals for Krebs Buffer

| Description | Source | Catalog # | Persistent ID / URL |
|---|---|---|---|
| NaCl | Sigma | 746398 | https://www.sigmaaldrich.com/US/en/product/sigald/746398 |
| D-glucose | Sigma | RDD016 | https://www.sigmaaldrich.com/US/en/product/sigma/rdd016 |
| KCl | Sigma | 746436 | https://www.sigmaaldrich.com/US/en/product/sigald/746436 |
| $NaHCO_3$ | Sigma | 792519 | https://www.sigmaaldrich.com/US/en/product/sigald/792519 |
| $CaCl_2 \cdot 2H_2O$ | Sigma | C5080 | https://www.sigmaaldrich.com/US/en/product/sial/c5080 |
| Na-HEPES | Sigma | H8651 | https://www.sigmaaldrich.com/US/en/product/sigma/h8651 |
| $MgSO_4$ | Sigma | 793612 | https://www.sigmaaldrich.com/US/en/product/sigald/793612 |
| $NaH_2PO_4 \cdot 2H_2O$ | Sigma | 71507 | https://www.sigmaaldrich.com/US/en/product/sigma/71507 |
| BSA | Sigma | A3311 | https://www.sigmaaldrich.com/US/en/product/sigma/a3311 |

### Figure 1 – Mice for scRNAseq Data

| Groups | Sex | Age | Number (prior to experiment) | Number (after termination) | Littermates (Yes/No) | Other Description |
|---|---|---|---|---|---|---|
| WT control | M | 3-6 months | 4 | 4 | Yes | n=14 vessels per mouse |
| WT control | F | 3-6 months | 4 | 4 | Yes | n=14 vessels per mouse |

### Figure 2 – Mice for Pressure Myography Data Semaglutide Dose Response Curve

| Groups | Sex | Age | Number (prior to experiment) | Number (after termination) | Littermates (Yes/No) | Other Description |
|---|---|---|---|---|---|---|
| WT control | M & F | 3-6 months | 10 | 10 | Yes | n=17 vessels total |

**Figure 3 – Mice for Pressure Myography Data Semaglutide Single Dose**

| Groups | Sex | Age | Number (prior to experiment) | Number (after termination) | Littermates (Yes/No) | Other Description |
|---|---|---|---|---|---|---|
| WT control | M & F | 3-6 months | 10 | 10 | Yes | n=11 vessels total |

**Figure 4 – Mice for Pressure Myography Data L-NAME+Indo Perfusion + Semaglutide Single Dose**

| Groups | Sex | Age | Number (prior to experiment) | Number (after termination) | Littermates (Yes/No) | Other Description |
|---|---|---|---|---|---|---|
| WT control | M & F | 3-6 months | 4 | 4 | Yes | n=11 vessels total |

**Figure 5 – Mice for Pressure Myography Data L-NAME+Indo+Apo Perfusion + Semaglutide Single Dose**

| Groups | Sex | Age | Number (prior to experiment) | Number (after termination) | Littermates (Yes/No) | Other Description |
|---|---|---|---|---|---|---|
| WT control | M & F | 3-6 months | 6 | 6 | Yes | n=11 vessels total |

**Figure 6 – Mice for Pressure Myography Data Semaglutide Perfusion**

| Groups | Sex | Age | Number (prior to experiment) | Number (after termination) | Littermates (Yes/No) | Other Description |
|---|---|---|---|---|---|---|
| WT control | M & F | 3-6 months | 7 | 7 | Yes | n=17 vessels control<br>n=10 vessels sema |
| DIO (western diet fed WT mice) | M & F | 3-6 months | 5 | 5 | Yes | n=13 vessels control<br>n=12 vessels sema |
| ApoE KO | M | 3-6 months | 7 | 7 | Yes | n=19 vessels control<br>n=9 vessels sema |
